# Diversity of Microarthropods in Different Habitats: An Ecological Perspective

**DOI:** 10.1101/668707

**Authors:** Mohammad Jalaluddin Abbas, Hina Parwez

## Abstract

Soil biological communities are influenced by several factors like edaphic factors, environmental variables, soil nutrients availability and anthropogenic changes including urban development and agricultural practices. *Edaphic factors exert the conditions directly to influence microarthropods populations*. To test this hypothesis, a question has been addressed; how factors regulate the ecosystem functions in contemporary a combined fashion? We observed, vegetation and habitat conditions both have close relationship and profoundly affect microarthropods populations particularly affect on local scale, resulted microclimatic influence of edaphic matrix which is generally system oriented. It is further discussed in this paper, on what attempt possible for better functional use of microarthropods which is important for conservation and for better output as a result of soil productivity in contemporary harsh climatic conditions with minimum use of amendments. It has been concluded in this study that diverse species found in structurally diverse habitats where they often appear to specialized conditions in native agricultural ecosystem.

## Introduction

Soil provides a great variety of microhabitats for its inhabitants from which soil microarthropods makeup significant proportion. These are different in size, generally 0.1-2 mm body width, (so called mesofauna, Swift et al. 1979), physiologically active and ecologically important indicators of soil conditions. Besides participating microarthropods in various soil ecological activities (decomposition of residues, mineralization of nutrients, improve the soil microstructure by their fecal palates, reduce the bacterial and pathogenic attack by consuming these organisms, and critically improve the porosity of soil by their vertical as well as horizontal movements), their species diversity facilitates maximum exploitation on the resource available in different soil habitats. Apart from soil microarthropods, Collembolans and Acarina (mites) might infer the degree of exploitation of available resources and complementarily affect ecosystem services. In most of Indian soil ecosystems, these two groups diversified up to 72-97 % among microarthropods (Reddy and Alfred 1992, Singh and Mukharji 1971, Singh and Pillai 1975, and Prabhoo 1976) thus dominate in most kind of soils (Zhu et al. 2010; Brahmam et al. 2010; Parwez and Abbas 2012). Therefore, there is an ample scope of use of these key biological components mostly in tropical and subtropical agricultural ecosystems.

Tropical and subtropical ecosystems exhibit specific ecological characteristics that make difficult situations for their sustainable management. Most preferentially in agricultural ecosystems, soil matrix and textural conditions, and food resource, all shaped by vegetation type; whereas the vegetation itself affected by all these factors including ecological factors. Therefore, rotation of vegetation is a general process of agriculture under crop cultivation. However, harsh climatic conditions such as drought, flooding and dry cold or warm conditions, dramatically affect the establishment of crop cultivation. In these conditions, soil inhabiting biological communities preferentially soil microarthropods faced survival problems in continuously changing edaphic environment. Thus, there is a need to study, how an ecologically changing environment affect microarthropods populations in an agricultural soil system? This can be evaluated by the study of ecological and edaphic factors in concerned study sites.

Previously several researchers have noted the effect of various ecological factors on distribution and abundance of microarthropods in different soil habitats. Most of these studies are on edaphic factors including, soil temperature (Usher 1976, Whitford 1989, Asikidis and Stamou 1991, Cancela Da Fonseca 1995, Sulkava and Huhta 2003, and, Cakir and Makineci 2013), soil moisture (Wallwork 1970, Usher 1976, Badejo 1982, Steinberger et al. 1984, Kamill et al. 1985, Vannier 1987, Whitford 1989, Asikidis and Stamou 1991, Bean et al. 1994, and, Ali-Shtayeh and Salahat 2010), soil pH (Hagvar and Abrahamsen 1980, Klausman 2006, and Rentao et al. 2013), soil organic matter (Fujikawa 1970, Santos et al. 1978, Anderson 1988, Scheu and Schulz 1996, and Ponce et al. 2011), and vegetation (Speight and Lawton 1976). These studies suggest that there is ample scope of investigation on edaphic factors in context of soil microarthropods to envisage the challenges of ecological manipulations due to which microarthropods abundance fluctuates in a highly varied manner.

### Problem formulation

There is currently an interest in understanding the factors which regulate the structure of soil faunal communities with consequences of ecosystem function (Cole et al. 2006). However, an interesting question is that, how factors regulate the ecosystem functions in contemporary a combined fashion? When considered the ecosystem function, it is likely depend on, how many species or communities are necessary in such ecosystem for proper functioning of an ecosystem? As we know, biological species live with competitive exclusion, without which they can’t survive in a more fascinate life. However, competitive exclusion may decline some species properly; because of the extent of varied situations and unavailability of suitable resources for survival critically affect on the diversity of communities. Therefore, existence of competition within soil microarthropods communities might be a contentious issue. The fact is that, each and every community has to be use maximum resources to survive and shape much diverse itself. On the other hand, soil biological communities are shaped by combined effect of several factors, like edaphic factors, environmental parameters, resource availability and anthropogenic influences, including urban development and agricultural practices. Apart from them, *Edaphic factors exert the conditions to influence microarthropods populations*. To test this hypothesis, we studied soil microarthropods populations in different native soil ecosystems.

## Materials and Methods

### (a) Area and sites of study

The area selected for study is situated at Aligarh. It is a flat topographical area, located in western part of UP, India at latitude 27-54’N, longitude 78-05E’ and altitude 187.45 meter above sea level. It is a subtropical zone with fluctuating climatic conditions consisting of four different seasons characterized by extreme winter and summer followed by medium to heavy rainfall during monsoon months and a post winter sweet spring. In hot dry summer, the temperature rises up to 48 °C sometimes near 50 °C, while in winter cold, the temperature goes drops up to 0-2 °C. Such widely varying climatic conditions provide a variety of ecological niche to soil dwelling animals and interesting for ecological studies on microarthropods in this region.

Four different study sites have been selected to collect the soil samples for the population study of soil microarthropods, namely Agriculture Quarsi Site (AQS), Agriculture Panjipur Site (APS), Mango Orchard Site (MOS) and Un Arable Site (UAS). AQS and APS were agriculturally well managed sites with the difference that AQS was totally organic managed site whereas APS was conventionally managed site. On the other hand, MOS was the orchard site which was less managed and UAS was unmanaged site without any human interference in field site.

### (b) Sampling, Extraction, Preservation and Identification of microarthropods

Samples have been taken from all study sites at regular weekly intervals in every month for two consecutive years (384 samples). Modified Tullgren’s funnel apparatus has been used for extraction of soil microarthropods. All microarthropods have collected inside a beaker which contained 70% ethyl alcohol with few drops of glycerol so that microarthropods are not getting dried. Microarthropods were separated and mounted directly in DPX. A binocular stereomicroscope (OLYMPUS, CX-21) has been used to identify soil microarthropods.

### (c) Measurement of Edaphic factors

The edaphic factors include soil temperature, relative humidity, and moisture content of the soil. Temperature of the soil has been measured by directly inserting the thermometer into the soil up to the required soil depth and relative humidity on soil surface was determined by Dial Hydrometer. The absolute content of water, which has an impact on the activities and distribution of soil microarthropods, exists in variable quantity in soil. Therefore, soil moisture has been measured by directly inserting digital moisture instrument up to the required soil depth. Soil pH was estimated by using digital soil pH meter. Organic carbon of soil was estimated according to Walkley (1934) method.

### (d) Statistical Analysis

We analyzed the correlation among the population of soil microarthropods and edaphic factors. SPSS version 20.0 has been used to carry out the statistical analysis. The lognormal (log10) basically used for cyclic differentiation of microarthropods population data (Figure 2a). Box and Whisker plot for microarthropods made by using Graf pad Prism 7.0 to show significant response and stability of populations. We also estimated population density of soil microarthropods (Abbas and Parwez 2009).

The indexes such as, Simpson Index, Shannon Index are evaluated as under-

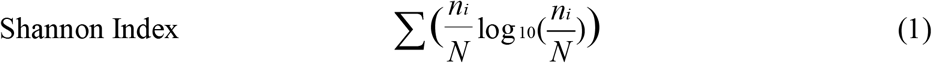

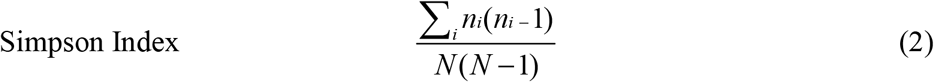

We examine the corresponding values of both indexes to check whether there is any relation between these two indexes or not if correlated.

## Results

Total 15,286 specimens of soil microarthropods have been collected from all four selected sites during investigation time. The population of microarthropods exhibit decreasing trend from first year (8,047 specimens) to second year (7,239 specimens). Seasonal variations of soil microarthropods exhibit different trends for different sites. It is observed that the density of soil microarthropods is high in winter and spring seasons at both the agricultural sites (AQS & APS) whereas in monsoon the density of soil microarthropods at Un-arable Site is relatively larger. Also, for Mongo Orchard Site, the density of soil microarthropods is observed to be relatively larger in summer and monsoon seasons (Figure1) than compare to rest of time.

The lognormal values of cyclic patterns of microarthropods populations have been the subject of remark for two consecutive years (Figure 2a). This confirms the population of microarthropods highly variable in order of AQS>APS>MOS>UAS. Generally single peak population found in both highly managed agricultural sites (AQS & APS) whereas bimodal peaks (Figure 1) are confirmed in rest two sites (MOS & UAS). On the other hand, both AQS and APS were highly abundant than compare to MOS & UAS (Figure 2b). However, seasonal rhythms are more effective in microarthropods population buildup than compare to temporal (monthly) variation in all the sites. The higher deviation of populations showed in both agricultural sites (Figure 2a) whereas lower populations but stability observed in Un Arable Site (Figure 2b).

**Figure 1.**
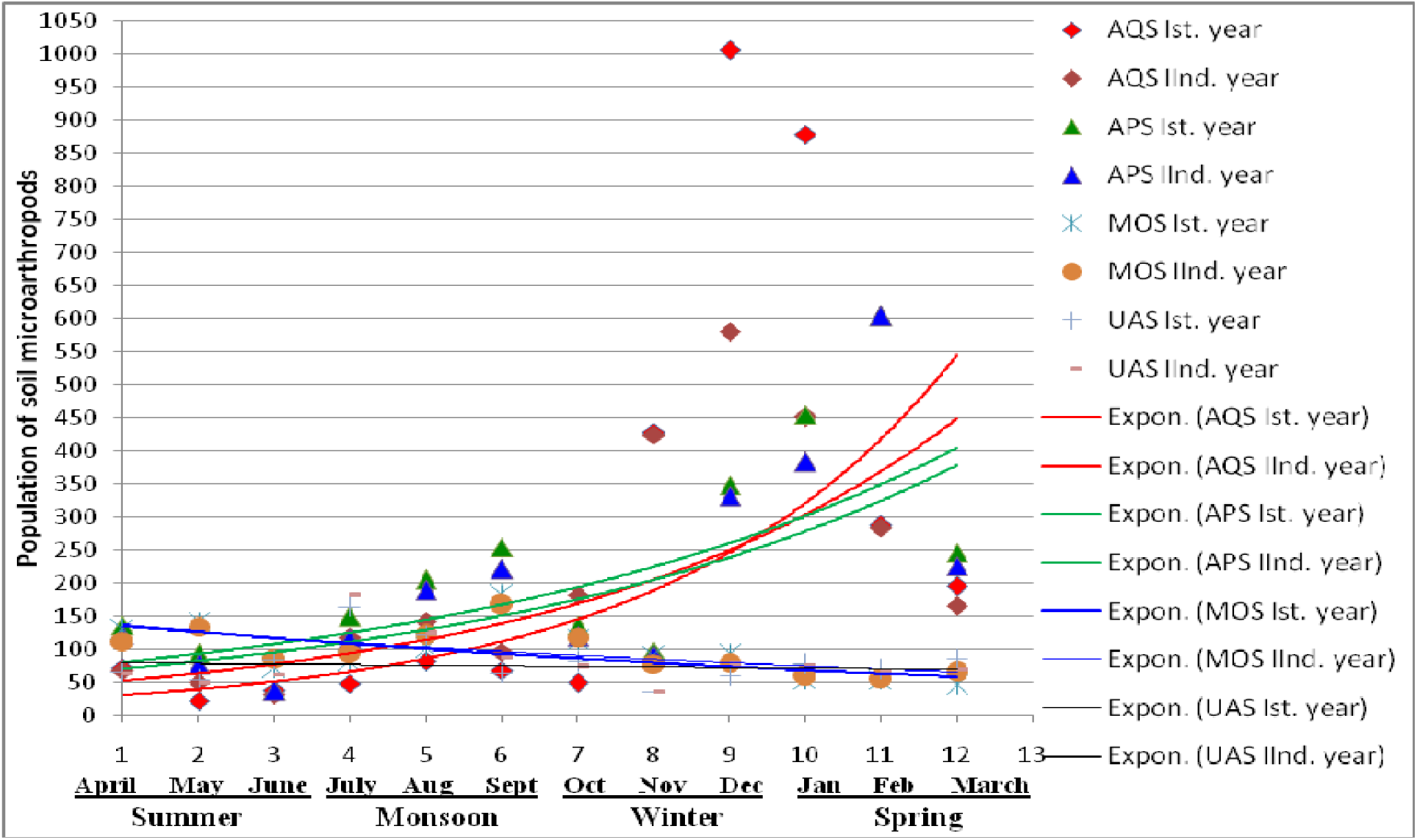
Variation of microarthropods in different study sites within two consecutive years.

**Figure 2.**
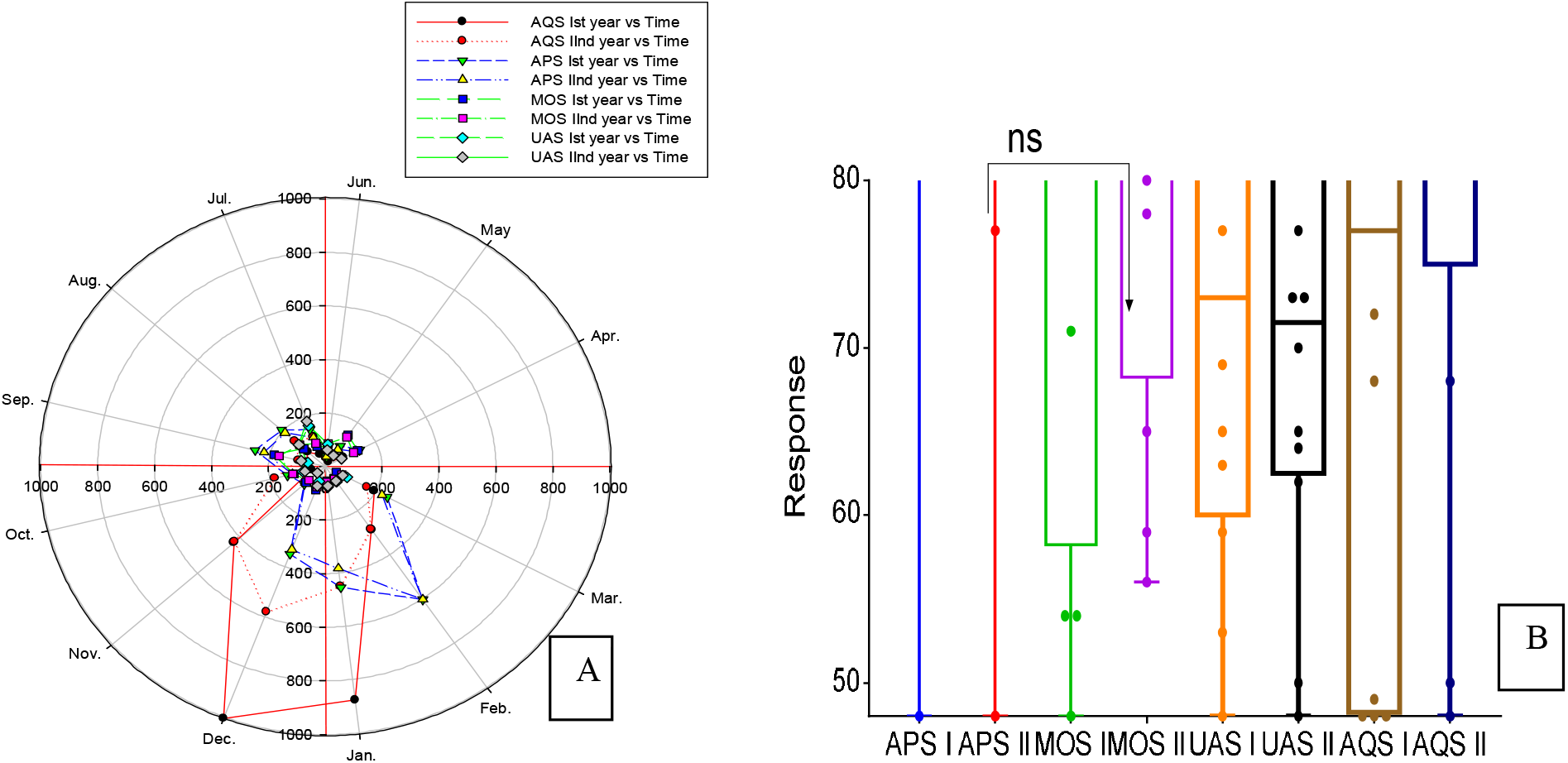
(A) Real limits of deviation of populations showed by polar plot (B) Box and Whisker plot for microarthropods populations’ response to significant using one way ANOVA.

### Relationships between soil microarthropods and soil matrix

In AQS, soil temperature always observed statistically negative significant (first year, p<0.05, □0.768 and second year, p<0.05, □0.573) whereas soil moisture always found statistically positive significant (first year, p>0.05, 0.736 and second year, p>0.05, 0.780) with reference to microarthropods population (Table 1). In APS, Soil temperature always found statistically negative significant with reference to the population of soil microarthropods as recoded in first year (p<0.01, □0.759) and in second year (p<0.01, □0.705). Soil organic carbon always found positively significant in all the sites except Mango Orchard site (Table 1).

**Table 1.** Correlation between edaphic factors and microarthropods populations.

| *Parameters* →<br>*Sites* ↓ | Density | Soil temperature (°C) | Soil moisture (%) | RH (%) | Soil pH | SOC (%) |
| --- | --- | --- | --- | --- | --- | --- |
| AQS Ist year | 66.08 | -0.768** | 0.736** | 0.282 | -0.640* | 0.786** |
| AQS IInd year | 54.02 | -0.573* | 0.780** | 0.425 | -0.470 | 0.491 |
| APS Ist year | 57.50 | -0.759** | 0.344 | 0.723** | -0.420 | 0.513* |
| APS IInd year | 52.42 | -0.705* | 0.302 | 0.718** | -0.422 | 0.605* |
| MOS Ist year | 19.96 | 0.439 | 0.261 | -0.075 | -0.394 | -0.168 |
| MOS IInd year | 20.02 | 0.578* | 0.221 | 0.052 | -0.837** | -0.245 |
| UAS Ist year | 24.10 | 0.391 | 0.246 | 0.344 | -0.300 | 0.697* |
| UAS IInd year | 24.35 | 0.183 | -0.054 | 0.394 | 0.094 | 0.359 |

Significant effects are given in red coloured values having \*\* Correlation is significant at $P < 0.01$ level and \* Correlation is significant at $P < 0.05$ level.

### Relationships between Shannon and Simpson’s diversity indices

We analyze two main diversity indices, Shannon and Simpson diversity index. Simpson’s diversity index found parallel that of Shannon’s diversity. Therefore, they are approximately corrective interdependent on each other as we recorded (Figure 3) in this experiment. This observation also introduces the differential between sites studied. The fact is that the higher populations always found in highly diverse habitat with diversity influence (Figure 3).

**Figure 3.**
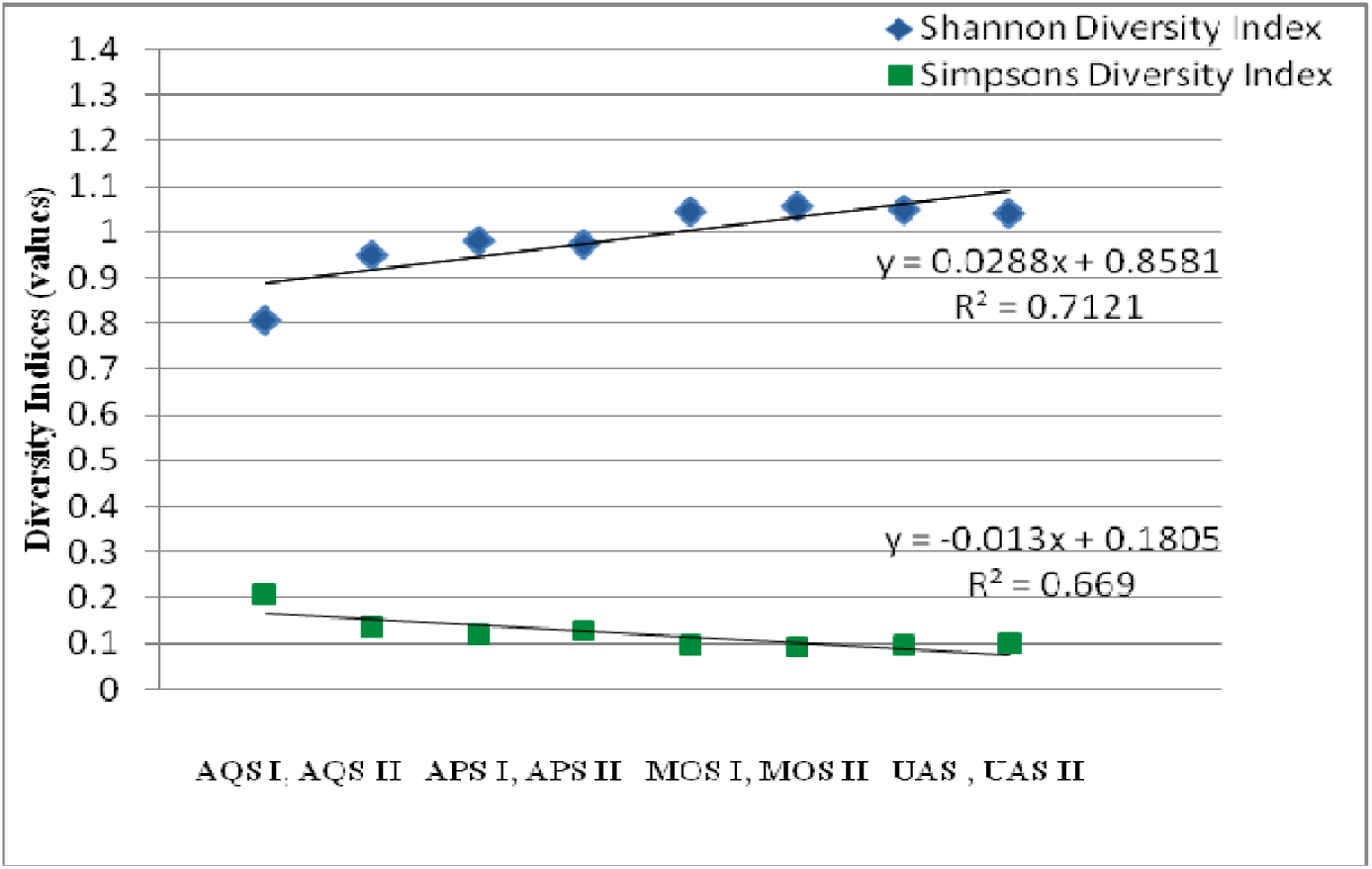
Comparative regression between Shannon and Simpson’s diversity

## Discussion

The close relationship of climatic variations, its affect on the species concern and resistibility of species affected, are inseparable (Parwez et al. 2011) in field conditions. Therefore, the alteration of soil faunal species depends on functional relationships of inter and intra climatic interferences. A large number of studies have documented that present global climatic change has significant consequences on flora, fauna and ecosystem processes worldwide (Graham & Grimm 1990, Walther et al. 2002, Parmesan & Yohe 2003, Weltzin et al. 2003), they may be temporal or seasonal. As observed in this study that, soil temperature is a critical factor in shaping the diversity of microarthropods in soil ecosystems; several researchers also reported that temperature is an important factor, regulates many aspects of their life (Christiansen 1964, Butcher et al. 1971, Hopkins 1997).

Changes in soil microarthropods populations associated with patterns of spatial and temporal variation might substantially due to changes in vegetation in an agriculture ecosystem because heterogeneity of soil microarthropods in an agricultural land is strongly correlated with the number of floral species or their density. Wardle et al. (2004), also stated same situations under ecosystems dominated with plant species adapted for fertile conditions can support high faunal densities, with more than 50% of net primary productivity (NPP) being returned to the soil as labile fecal material. Fertile conditions are therefore likely to support rapid, leaky nutrient cycles and low net accumulation of soil carbon, whereas infertile conditions support slow nutrient cycles and soil carbon sequestration is promoted (Wardle 2002, Coleman et al. 1983) which is less imparting in growth and support of microarthropods; ultimately meager population of soil microarthropods are observed (both in UAS & MOS). This is much clear from the observations of this study, because most of the populations of soil microarthropods were highly abundant during dense floral vegetation or crop (i.e wheat crop in both agricultural sites during mid October to last March, Figure 1) in both agriculturally managed sites compare to less managed mango and un-arable site.

Agro-ecosystems differ from natural (un-arable) ecosystems in terms of management, soil biodiversity and nutrients output via crop harvest for exceed nutrients loss pathways in combined sequences because, agriculture replace the nutrients lost via crop harvests and many amendments are also added either inorganic or organic fertilizers. Aside from amendments added, organic amendments improve soil physical properties and help maintain soil organic matter contents. However, the effect of organic inputs on physical properties of soil depend on the time residence, quality and placement of the material concerned, means that, the longer the residence, the more durable the effect, might intern the higher quality of organic carbon in soil. Thus, population of soil microarthropods found higher in managed sites than compare to un-arable (non-managed) and less managed sites as observed in this study (Figure 1).

It is clear from the growing number of ecological network studies that, biotic interactions often take place in multispecies frameworks, rather than few species-specific pair wise interactions (Stanton 2003). Thus, it has been widely recognized that the interactions between species, such as predator–prey relationships or competition within a guild, are crucial for shaping community dynamics (May 1974). Rapid increase of a community might be a surprise to those who expect that organisms form permanent mutualistic associations more likely to be highly specific, ensure a result of competitive exclusion.

Gause (1934) stated in competitive exclusion theory that differences in competitive abilities can cause non-random patterns of species occurrences among sites and generate inequalities in species abundances within sites. As per another study cleared that, competitively inferior species are predicted to occur less frequently and at lower abundance which has an important and largely unresolved question; how such species can persist in a community over long time periods (Fox 2013). Actually, this depends upon less frequent environmental conditions favorable for particular species. The reason is that, competition can moderate the species concerned, which may be depend on environmental heterogeneity (Allesina and Levine 2011). Thus, environmental heterogeneity increases the competition between species and within the communities. In case of soil microarthropods, it is much clear that they are much more sensitive to the environmental variations. Shakir and Ahmad (2014) recently stated that, soil arthropods respond very quickly to the changes in the environment. Recently, Tsiafouli et al. (2015) stated that, not only diversity of many taxonomic groups decreased and communities were composed of more closely related species, but also some functional groups may be completely lost under intensive agriculture with potential implications for ecosystem functioning.

Phelan’s (2009) statement was that an important function of ecosystem is community stability. Phelan addressed community stability expressed in relation to three parameters (Barbour et al. 1999 also confirmed); (1) resistance—the ability to remain unchanged during a period of stress, (2) resilience—the ability to return to its original state following stress or disturbance, and (3) persistence—the ability to remain relatively unchanged over time., We largely determined by 1 and 2 among these parameters. The aggregations of individual organisms is actually depend upon the favorability of environment either climatic or field conditions rather than made up of organisms whose characteristics are complement to each other and functions are highly integrated. However, presence or absence of other organisms affects a species ability to exploit or tolerate conditions, not only through competition or mutualism, but also through their impact on the habitat itself, either opening up new niches or shrinking niche dimensions for other species (Sterelny 2005). The cumulative effect of the biotic components of an ecosystem affects the local environment in a nonlinear fashion (Phelan 2009). However, a saturated community represents a state of equilibrium, governed largely by competitive interactions between species (Giller 1996) and within species. Therefore, stability tends to greater (UAS) in soil ecosystems where a larger number of functionally diverse species were present (Figure 2a & 2b).

On the other hand, habitat has abrupt effect on the life cycle of microarthropods and enhances their population diversity (Dowdy 1965). The interaction between drought and soil carbon enrichment may also be the consequent measure which directly affect performance of concerned soil ecosystem. This is due to that soil moisture content strongly controls carbon availability in soils. The pressure of influential temperature also reduces when soil has suitable and stable moisture conditions. In this investigation, more stable soil moisture conditions found in AQS and APS sites, favour the higher concentrations of available carbon contents in soils of both study sites which ultimately influenced the populations of soil microarthropods.

Soil microarthropods are positively recorded with soil moisture although these correlations largely depend upon season and site sampled (Kamill et al. 1985; Ali-Shtayeh and Salahat (2010). These results coincide with that of Wallwork (1970); Usher (1976); Vannier (1987); Whitford (1989); Asikidis and Stamou (1991). Soil pH was negatively correlated with soil microarthropods abundance although the correlation was sometimes weak (Table 1). Similar observation was reported by Klausman (2006) who found negative correlations between soil pH and microarthropods. Overall, changes in edaphic matrix affect the soil characteristics which in turn influence the soil microarthropods abundance. Thus, it is confirmed that, *edaphic factors exert the conditions to influence population of soil microarthropods*.

Vegetation types tend to occur along with the gradient of available moisture in soil. Available moisture was used here as the amount of moisture available to the plants (agricultural sites) that is necessary for their establishment and maintenance. This may be the reason by which higher population found in agricultural sites than compare to Un-arable site. Moisture gradient can be steep or gradual. Consequently, an interesting feature of gradual increase of moisture contents in soil directly increase influence the density of soil microarthropods. Soil moisture levels declined rapidly when temperature of soil increases. Moisture contents of soil underneath the irregularly varied between seasons with least in summer and highest in Rainfall. Therefore, the population of soil microarthropods was susceptible recorded in these periods (Figure 1). Hence, it may suggest that stability of edaphic environment specifically soil moisture with less fluctuation of soil temperature can lead the survival of soil microarthropods.

Vegetation also protects soil surface to warm from direct solar radiation/sun light rays therefore soil temperature found lower in agriculturally managed sites (AQS & APS) mostly in dry period (summer). Oppose to this indication, vegetation keep warm the soil surface in winter by avoiding the cool air due to abundant crop plants. The cold interference of climate induced variations due to which better survival conditions adopted by microarthropods in winter which cleared and observed in both agricultural management sites (AQS & APS). In agricultural lands (AQS & APS), perennial vegetation has been setup for the purpose of farming. It has been reported that these sites were highly diversified than compare to other two less managed (MOS) and unmanaged UAS. The reason may be that, increase root microbial diversity and their activities enhance nutrient cycling and possible allocation of more carbon into deeper horizons of soils (Millard and Singh 2009). Furthermore, perennial vegetation strips within agricultural fields may accumulate soil organic carbon relatively quickly because of runoff retardation, improved hydraulic properties, and nutrient deposition in the vegetative buffer strips (Udawatta et al. 2002, 2008; Udawatta and Jose 2011) compare to plantations in orchards or in un-arable fields. This may be the strong reason for higher population buildup of soil microarthropods as well as higher soil carbon contents observed in both agricultural sites (AQS & APS-Table 1) compare to other two sites (MOS & UAS-Table 1).

Climate change in few past years causes a seasonal shift of monsoon in Indian continent mostly in North Central part of India, resulting in less frequent however, more extreme events during summer (resulted increased temperature up to 48 °C) as well as in winter (resulted declining temperature up to 0 °C). This may prognosticate much-more changes in total annual rainfall in future. The direction, magnitude and variability of such changes in precipitation events and their effect on soil systems functioning will depend on how much the change deviates from the existing variability and the ability of soil ecosystems and their inhabiting soil organisms to adapt to new conditions (Beier et al. 2012). Thus, economically some important aspects be summarized as- (i) adaptability or response of soil biological constraints might shape the functional potential of an ecosystem; (ii) increasing uncertainty of environmental changes profoundly affect soil biological constraints; and (iii) environmentally site specific effect may intern the survival of microarthropods resulting decline of some species up to extinct situations in an ecosystem.

## Conclusion

The sustainability of agricultural ecosystems and their responses depend on the continuation of essential ecological processes. Ecological processes are depend on the type and intensity of management within particular landsite for what purpose to be achieved in one hand and seasonal manipulations in other hand, in tropical and sub tropical countries. Extreme events of seasonal patterns might disrupt the biological populations. However, the management can serve the ecological communities by enhancing the active ecological processes. Ecological processes depend on what functional contribution of habitat specific for its inhabitants. Therefore, continuous and reliable delivery of ecosystem functions, depend on the maintenance of functionally-diverse ecological assemblages.

